# A Virtual Reality Dataset to Support Hand Action Observation in Rehabilitation and Motor Learning Studies

**DOI:** 10.64898/2026.02.23.707557

**Authors:** A. Ciraolo, E. Scalona, A. Zilli, A. Nuara, D. De Marco, G. Rizzolatti, P. Adamo, R. Gatti, P Rossi, S Banfi, M.A. Rocca, M. Filippi, P. Avanzini, M. Fabbri-Destro

## Abstract

The recovery of motor function is increasingly understood as a process influenced not only by physical training but also by perceptual and cognitive strategies. Action Observation Treatment (AOT), a neurorehabilitation approach in which patients observe goal-directed motor actions before executing them, has demonstrated clinical benefits; however, its wider implementation is hindered by a lack of standardized procedures. We present an open-access dataset of 33 upper limb gestures specifically developed to support the administration of Virtual Reality-based AOT (VR-AOT). The gestures were selected in collaboration with expert physiotherapists to ensure clinical relevance, and are provided as motion capture recordings along with Unity-based 3D animations embedded in configurable virtual scenes. The dataset is designed for flexibility, allowing users to modify parameters such as viewpoint, laterality, and repetition count. Technical validation confirms its usability and therapeutic applicability across multiple clinical and research contexts. This dataset offers a standardized yet customizable resource for developing and comparing VR-AOT protocols, with potential applications in neurorehabilitation and motor learning research.

## Background & Summary

Over the past few decades, advances in cognitive neuroscience, neuroimaging, and neurophysiology have broadened the range of strategies available to guide, support, and enhance rehabilitation. These approaches now extend beyond traditional muscle-and joint-based interventions to include central strategies such as action observation^1,2^, motor imagery^3,4^, and non-invasive brain stimulation. Among them, Action Observation Treatment (AOT) has emerged as particularly promising. AOT leverages the human mirror mechanism, i.e., the neural principle whereby the observer’s motor system is activated in a manner corresponding to the execution of the same action^5,6^. This motor resonance is not limited to the neuronal level but also extends up to corticospinal excitability^7^, inducing facilitation whose time course mirrors the unfolding of the observed action^8^. These features make action observation an invaluable component in processes aimed at promoting motor learning or recovery, given its potential to pre-activate the motor system in an externally driven manner before actual execution. Building on this principle, AOT typically chains action observation with motor imagery^9^ (see for a meta-analysis documenting the neural substrates of these processes, along with their overlap), before engaging the individual in physically performing the action.

This protocol has shown significant potential for enhancing motor learning in healthy individuals^10,11^, limiting motor deterioration following limb immobilization^12^, and supporting recovery in both adult and pediatric populations across various clinical conditions^2^.

Despite its robust neurophysiological foundation, AOT faces several challenges that hinder its broader implementation and optimal efficacy. One key limitation is the lack of standardized stimuli and procedures^2,13^. Existing AOT protocols vary widely in terms of action types and gesture complexity, with the stimuli selected, videotaped, and administered by the clinical personnel responsible for the patient’s rehabilitation protocol. This results in substantial variability across studies, which limits replicability, reduces clinical scalability, and complicates comparisons. These issues highlight the need for open-access, standardized AOT stimulus repositories.

Beyond action type, recent research indicates that various stimulus features—such as viewpoint^14,15^, number of repetitions, and the use of 3D visualization^16^—modulate neural responses to action observation and may influence the effectiveness of AOT. While traditional video-based stimuli offer limited flexibility, virtual reality (VR) provides a promising alternative. VR’s immersive and interactive nature enables the delivery of more naturalistic and controllable action stimuli, allowing for the precise manipulation of parameters such as perspective, timing, and movement complexity. Preliminary findings suggest that VR-based AOT (VR-AOT) may enhance patient engagement and increase activation within the action observation network^12,17^.

However, integrating VR into AOT remains logistically challenging. Developing VR-based stimuli typically requires multidisciplinary expertise, including software engineering, 3D animation, and programming, as well as access to specialized hardware such as head-mounted displays. These demands limit accessibility, particularly in clinical and resource-constrained environments, making the creation of public VR-AOT repositories even more fundamental for the adoption and diffusion of such a training strategy. Currently, there is no comprehensive VR-based repository of AOT stimuli. Existing resources are scarce and either focus on videotaped action observation^18^ or target VR stimuli, yet in highly specific contexts such as occupational therapy^19^.

To address these gaps, we present a freely accessible and standardized dataset comprising 33 upper limb gestures optimized for VR-based AOT. The dataset is accompanied by a Unity project that integrates the animations into a customizable virtual reality (VR) platform. This system allows users to adjust key parameters such as the number of repetitions, viewpoint, and limb laterality, making it adaptable to a wide range of research and clinical scenarios. Such flexibility is particularly valuable in rehabilitation settings, where individual variability and contextual demands require tailored interventions. The repository is available both as a Unity package and as a precompiled Android application, improving accessibility for users with limited technical expertise. This dual format enhances usability while maintaining scientific rigor, supporting current efforts to translate neurotechnology innovations into real-world therapeutic applications^20,21^.

By offering high-quality, standardized stimuli and a flexible VR platform, this resource aims to facilitate the development, testing, and personalization of AOT protocols in experimental and clinical settings. It promotes reproducibility, enables more robust cross-study comparisons, and provides a foundation for developing individualized, technology-supported training programs.

## Methods

In this section, we describe how we created our VR dataset starting form kinematic data.

### Data Collection

Whole-body kinematic data were acquired from a 24-year-old, right-handed healthy volunteer^22^, during the execution of 33 exercises commonly used in upper-limb motor rehabilitation. Procedures have been approved by the CNR Research Ethics and Integrity Committee (project QUASAR, 22 September 2022), and informed consent was obtained from the volunteer. The exercise set was identified and selected in collaboration with expert physiotherapists from the Humanitas Clinical and Research Center (Milan, Italy), who also supervised the acquisition sessions to ensure clinical appropriateness and procedural accuracy.

Kinematic data were recorded using a dual-system setup. The primary motion capture system was the Xsens Biomech Awinda (Xsens Technologies, Netherlands), which features 17 miniature inertial measurement units (IMUs) that integrate 3D accelerometers, gyroscopes, and magnetometers. These sensors were placed on the participant’s body using manufacturer-specified straps, enabling the tracking of 23 body joints and segments, including:

- Head and neck
- Eighth and tenth thoracic vertebrae
- Third and fifth lumbar vertebrae
- Right and left shoulder, arm, forearm, and hand
- Pelvis
- Right and left thigh, shank, foot, and forefoot

To obtain a clearer picture of hand and finger kinematics, Manus Prime II Xsens gloves (Manus, Netherlands) were used in conjunction with the Awinda system. Each glove features industrial-grade flex sensors and integrated inertial measurement units (IMUs), enabling precise tracking of three joints per finger.

Before recording, anthropometric measurements of the subject were obtained to allow scaling of the biomechanical model. Then both static and dynamic calibration was performed to enable the alignment of the reference systems of the inertial sensors with those of the body segments.

### Data Integration in Unity

The 33 gestures performed by the actor (see next paragraph) have been transformed into 33 VR-based animations, in line with a previous study from our group on the rehabilitation of occupational gestures.^19^

Kinematic data were processed using Xsens MVN Analyze software to compute the orientation and position for each body segment and each finger. The resulting skeleton-based animation data were exported in FBX format, compatible with Unity 3D for real-time animation.

The FBX files containing the motion capture data were imported into the Unity 3D game engine software (version 6000.0.24f1) to animate a rig humanoid avatar (available at https://renderpeople.com/free-3d-people). For each exercise, colliders were placed on the avatar’s hand and the objects that had to be manipulated. This ensured the realistic rendering of hand-object interactions. When a collision occurred between the hand and an object, the object’s position was dynamically aligned with the finger kinematics until it was released. The object release was implemented through delegated animation events, by marking the exact frames in the animation where hand-object contact was lost. Finally, each animation was reviewed for consistency checks and, if necessary, edited using the Unity animation tool to adjust body segment postures or movements. Finally, given that the original recordings were performed using the right arm, the animations were mirrored within Unity to produce equivalent left-arm gestures.

**Figure 1.**
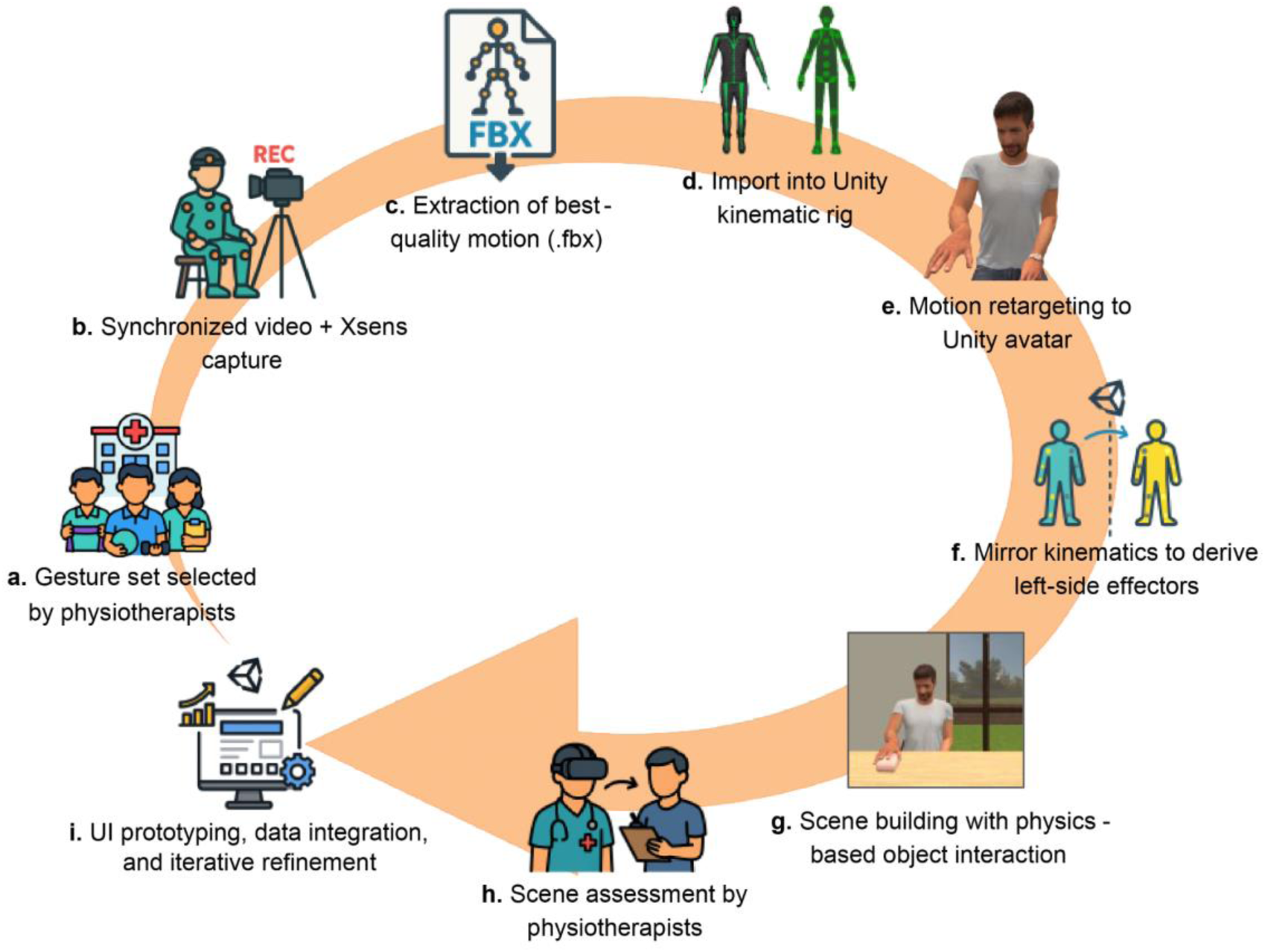
Workflow for generation and validation of upper-limb VR-AOT stimuli. a) Selection of a core set of upper-limb gestures by physiotherapists based on clinical frequency and relevance; b) Synchronous recording of each gesture using RGB video and the Xsens MVN Awinda inertial system (17 wireless IMUs, 60 Hz); c) Export of the highest-quality recordings in *.fbx* format for compatibility with the Unity animation pipeline; d) Import and mapping of .fbx files to Unity’s native kinematic rig; e) Retargeting of motion data to a standard Unity humanoid avatar using the built-in rigging system; f) Kinematic mirroring of right-arm stimuli to generate corresponding left-arm ones; g) Assembly of a virtual environment with physics colliders, inverse kinematics, and custom assets to enable realistic hand-object interaction; h) Evaluation of all scenes by 42 expert physiotherapists (see *Technical Validation section*); i) Integration of results into the Unity project to refine the user interface, codebase, and final dataset.

### Scene Description

Table 1 provides a comprehensive list of the 33 upper-limb exercises, designed for unimanual performance in a seated position. The task complexity ranges from basic proximal joint movements (e.g., shoulder flexion) to advanced fine motor activities (e.g., finger flexion). This progression enables therapists to tailor exercises to various stages of recovery. For example, the early rehabilitation phase may emphasize gross motor tasks that target the shoulder and elbow, while more advanced stages can incorporate precision tasks involving the wrist and fingers, according to the patient’s progression. This modular design supports the creation of personalized, adaptive rehabilitation plans tailored to patients’ abilities and clinical goals. To visualise how the graded tasks are instantiated in the virtual environment, Figure 2 displays representative frames captured from two exercises, each rendered in both allocentric and egocentric perspectives.

**Figure 2.**
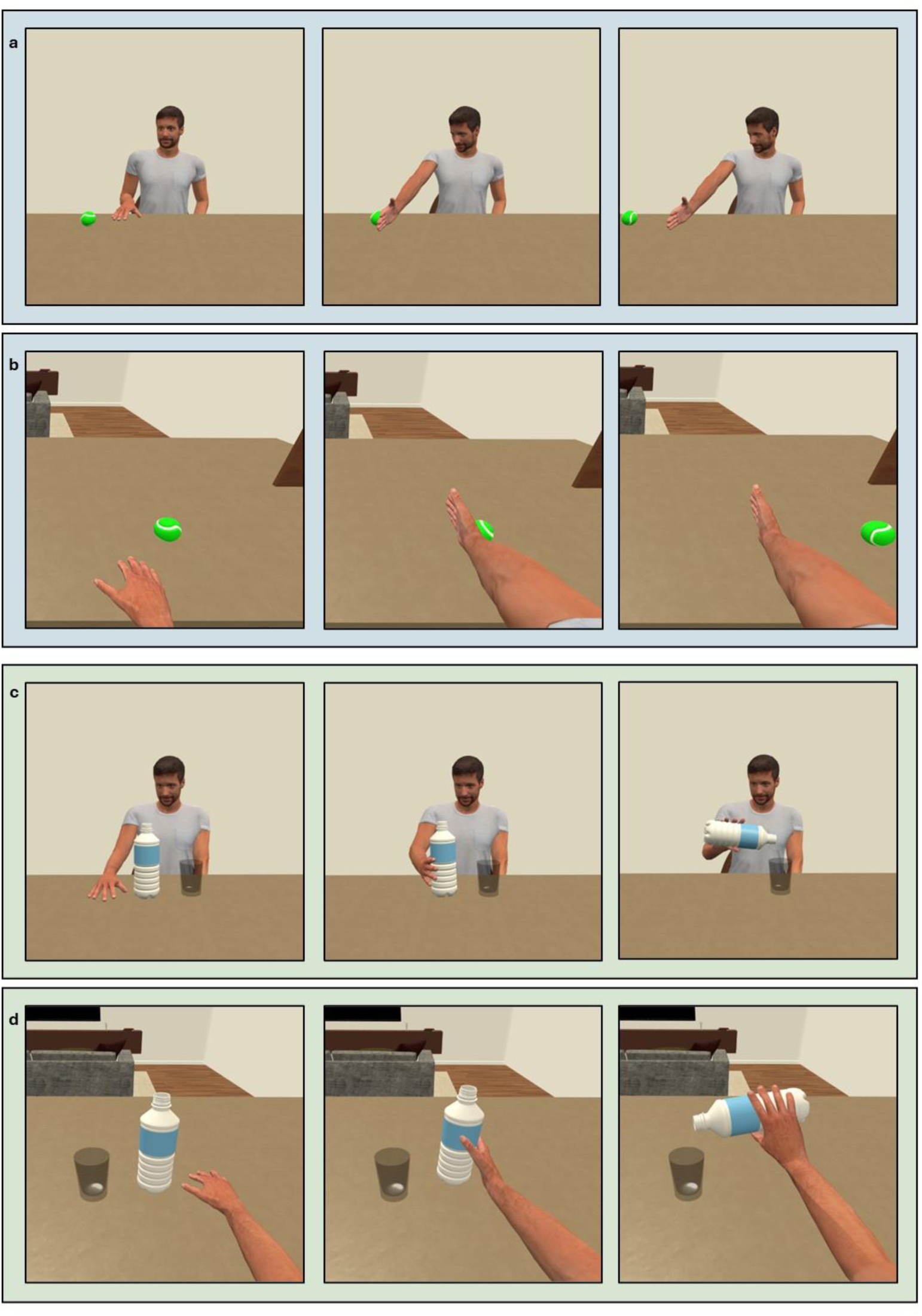
Temporal and perspective-based visualization of two representative VR-AOT actions. Representative frames from two actions shown across three temporal phases: rest (left), interaction onset (center), and completion/execution (right). The top two rows (blue background) depict a simple gesture (i.e., striking a ball with the back of the hand) from allocentric (a) and egocentric (b) perspectives. The bottom two rows (light green background) show a complex, goal-directed action (i.e., grasping a bottle and pouring water into a glass) in allocentric (c) and egocentric (d) views.

**Table 1.** Upper-Limb Rehabilitation Exercises: Scene Names, Movement Descriptions, and Involved Joints. The table outlines 33 goal-oriented exercises (E1–E33), each implemented as an independent Unity scene. The *Scene Name* column corresponds to the identifier used within the application; *Movement Description* provides a summary of the requested motor task.

|  | Scene Name | Movement Description |
| --- | --- | --- |
| E1 | Hit Ball - Dorsal | Hit a ball on a table using the back of the hand, keeping the elbow extended. |
| E2 | Hit Ball - Palmar | Hit a ball on a table using the palm of the hand, keeping the elbow extended. |
| E3 | Hit Ball - Flex | Hit a ball on a table by flexing the elbow. |
| E4 | Hit Ball - Ext | Hit a ball on a table by extending the elbow. |
| E5 | Drag Soap | Drag a soap on a table using the back of the hand with the elbow extended. |
| E6 | Drag Ball | Drag a ball on a table using the back of the hand with the elbow extended. |
| E7 | Drag Cloth - Flex | Move a cloth on a table using an elbow flexion-extension motion, with the forearm resting on the table. |
| E8 | Reaching Knee-Table | Reach a target placed on a table, starting with the hands on the knees. |
| E9 | Reaching High Target | Reach a target positioned above table level, starting with hands on the knees. |
| E10 | Reaching Lateral | Starting with the arm along the body, move and place the hand on the lateral surface of the contralateral thigh. |
| E11 | Drag Bottle | Move a bottle on the table to the point of maximum elbow extension. |
| E12 | Move Can | Reach for, grasp, and place a can on a surface 10 cm high. |
| E13 | Bring Soap | Reach for, grasp, and lift a bar of soap using elbow flexion. |
| E14 | Open Box | Reach, grasp the top of a box, and open it using elbow flexion/extension and forearm rotation (pronosupination). |
| E15 | Drag Box | Move a box on a table first to the right and then to the left. |
| E16 | Drag Journal | Move a journal on the table until full elbow extension is reached. |
| E17 | Pour Bottle | Reach, grasp a bottle, and pour water using elbow flexion/extension and pronosupination. |
| E18 | Clean Table | Clean a table using circular shoulder movements with a cloth. |
| E19 | Rolling Pin | Roll a rolling pin on a table using the palm. |
| E20 | Flip Glass | Grasp and flip over a glass. |
| E21 | Press Phone-Key | Press keys on a telephone keypad using the index finger. |
| E22 | Play Piano | Play adjacent piano keys using all five fingers, with the elbow resting on the table. |
| E23 | Hammer | Use a hammer to strike a nail on a table with elbow flexion-extension. |
| E24 | Round Spoon | Hold a teaspoon and rotate it along the edges of a cup using wrist movement. |
| E25 | Plug-Unplug | Insert and remove a plug from a tabletop socket using elbow flexion and extension. |
| E26 | Grasp Cubes | Grasp small cubes with the first three fingers and place them into an adjacent container. |
| E27 | Open Jar | Screw and unscrew the lid of a jar. |
| E28 | Handle | Rotate a tabletop handle. |
| E29 | Writing | Draw a circle on paper with a pen. |
| E30 | Sprayer | Pull the trigger of a spray bottle. |
| E31 | Throw Ball - L | Throw a tennis ball upward and forward, starting with the arm at the side. |
| E32 | Throw Ball - H | Throw a tennis ball downward and forward, starting with the arm at the side. |
| E33 | Tooth Brush | Simulate brushing teeth with a toothbrush. |

### Data Records

The complete dataset is available through Zenodo [https://doi.org/10.5281/zenodo.15879945] and comprises five primary components:

1. **UpperLimbAOT-VR_UnityProject/** — a fully self- contained Unity 6000.0.24f1 project directory.
2. **UpperLimbAOT-VR.apk** — a precompiled Android application package for immediate deployment on Meta Quest- series head- mounted displays.
3. **QuickStart_Guide.pdf** — a step- by- step installation and usage manual detailing project setup on both Unity Editor and Meta Quest devices.
4. **README.md** — a brief project overview and fundamental usage notes.
5. **Videos/** — a supplementary folder containing video demonstrations of the actions. This folder is divided into **AllocentricView/** and **EgocentricView/**, each of which includes **Left/** and **Right/** subfolders containing .mp4 files that show the recorded actions from both perspectives.

### Project root and directory convention

All domain-specific materials are nested beneath “Assets/UpperLimbAOT-VR/”, introducing an intermediate namespace (*UpperLimbAOT-VR*) that segregates experimental assets from Unity- generated infrastructure (e.g., Assets/Editor/ and Packages/) and thereby streamlines version control.

### Scene inventory

All scenes are located within **Assets/UpperLimbAOT-VR/Scenes/**.

- **ControlScene/Control. Unity** — the principal interface responsible for initial configuration and for orchestrating stimulus presentation (ordering, difficulty, viewpoint, repetition).
- **Allocentric perspective**

– AllocentricView/Right/ — 33 third- person (allocentric) scenes depicting right- arm movements.
– AllocentricView/Left/ — 33 third- person scenes depicting left- arm movements.
- **Egocentric perspective**

– EgocentricView/Right/ — 33 first- person (egocentric) scenes depicting right- arm movements.
– EgocentricView/Left/ — 33 first- person scenes depicting left- arm movements.

Lighting prefabs are co- located with each scene to ensure photometric consistency across perspectives.

### Supporting assets

All supporting assets listed below are located within Assets/UpperLimbAOT-VR/.

- **3DModels/** — static meshes and rigged avatars, including the humanoid skeleton and environmental props utilised across the scenes.
- **GraphicAssets/** — raster textures, user-interface sprites (.png, .tga), URP Asset renderer settings, skybox configurations, and post-processing data utilised within the project’s rendering pipeline and user interface components.
- **Kinematics/** — authorial motion data: the animator state machine in Controller/, together with limb- specific animation clips within Left/ and Right/.
- **Materials/** — includes physically based shader materials utilised within Unity’s Vulkan rendering pipeline, as well as material assets associated with specific 3D models (e.g., cups, boxes, nails) employed in the experimental scenes.
- **Prefabs/** — reusable GameObject assemblies comprising selected scene objects and various interaction widgets utilised throughout the project.
- **Scripts/** — bespoke C# scripts encompassing the scene- loading routine, randomisation controller, and data- logging subsystem.
- **StreamingAssets/** — high- definition instructional videos (.mp4, 1920 × 1080 @ 30 fps, H.264) streamed during runtime by the control scene. **Note:** for correct loading in Unity, this folder must reside directly at Assets/StreamingAssets, *not* inside Assets/UpperLimbAOT-VR/.

This arrangement provides researchers, clinicians, and software developers with a clear, modular structure that facilitates the rapid replication, adaptation, and extension of experimental stimuli.

### Technical Validation

In this section, we validate the clinical relevance and perceptual validity of the VR stimuli administering them to 42 physiotherapists (28 female, 14 male), with a mean age of 40.6 years (SD = 10.4; median = 36.5) and 12.3 years of clinical experience (SD = 6.8, median = 10). Regarding prior knowledge, 71% of participants were familiar with AOT (χ² = 7.71, *p* < 0.05), and 40% had previously administered AOT in clinical practice (χ² = 1.52, *p* = 0.217). In contrast, the majority had limited exposure to VR technologies: 79% reported no prior VR experience (χ² = 13.71, *p* < 0.001), and 95% had never used VR in combination with AOT (χ² = 34.38, *p* < 0.001).

### Design of the validation questionnaire

Each participant reviewed the full set of 33 stimuli and completed a questionnaire (available in the data repository), which was designed to evaluate multiple dimensions of the stimuli. Procedures regarding the validation questionnaire have been approved by the Research Ethics Board (REB) of the University of Parma (ID 23-2024-N, 28 February 2024), and signed informed consent was obtained from each participant.

Participants rated each stimulus across the following criteria:

1. *Comprehensibility:* Clarity of the gesture and ease of understanding the task;
2. *Realism*: Perceived accuracy and fidelity of the movements;
3. *Immersivity*: Degree of experiential engagement and immersion;
4. *Egocentric Embodiment*: Sense of ownership when observing from a first-person viewpoint;
5. *Allocentric Embodiment:* Sense of ownership when observing from a third-person perspective.

In addition to perceptual ratings, participants assessed the clinical applicability of each gesture for:

- The six stages of stroke recovery as defined by the Brunnström^23^ (stages I–VI).
- Central neurological disorders (e.g., Stroke, Multiple Sclerosis, Parkinson’s Disease, Pediatric Cerebral Palsy, and Traumatic Brain Injury);
- Peripheral conditions (e.g., Peripheral Nervous System pathologies, orthopedic trauma, and prosthetic rehabilitation).

All stimuli were rated on a 5-point Likert scale (1 = strongly disagree to 5 = strongly agree), with a score of 3 representing the neutral threshold that separates negative scores (<3) from positive ones (>3). Before proceeding with the statistical analyses, a Shapiro-Wilk test was applied to each distribution to verify the assumption of sphericity. Given that most of these tests failed, we adopted non-parametric approaches for the subsequent analyses. Wilcoxon signed-rank tests were performed to assess whether ratings for Comprehensibility, Realism, and Immersivity significantly exceeded the threshold of 3. In contrast, differences between Egocentric and Allocentric Embodiment ratings were analysed using paired-sample t-tests. Given that the same test was applied to 33 exercises, the significance threshold was adjusted using the Bonferroni correction. In addition, the interdependence between Comprehensibility, Realism, and Immersivity scores was tested with a non-parametric Spearman correlation.

Since one of the major applications of AOT concerns the upper limb rehabilitation of stroke patients, additional analyses assessed the utility of gestures across the six Brunnström stages. Wilcoxon signed-rank tests against a three-mean distribution (*p* < 0.05, Bonferroni-corrected) were conducted for each stage. Additionally, Spearman’s rank correlation was tested between the perceived difficulty and clinical utility of each gesture across the recovery stages.

A similar approach was applied to assess the perceived utility of the stimuli for other clinical conditions.

### Perceptual quality of stimuli

The large majority of the exercises were rated positively (scores significantly greater than 3) on Comprehensibility (average 4.5), Realism (average 3.8), and Immersivity (average 4.4) (Table 2), confirming that the stimuli were interpretable and biomechanically plausible. The highest comprehensibility scores were recorded for six exercises *(Hit Ball - Flex, Drag Cloth– Flex, Drag Bottle, Pour Bottle, Clean Table, Sprayer;* means = 4.8), accompanied by equally positive and significant immersivity ratings. While most gestures received strong realism ratings, a small subset (e.g., *Open Jar*, *Throw Ball – H*, and *ToothBrush*) yielded intermediate scores (Realism ≤ 3.5), indicating that some critical aspect of the rendering of the gestures was not fully captured. The three scores resulted positively and significantly correlated with each other (Realism vs Comprehensibility: Rho=0.84; Realism vs Immersivity: Rho=0.82; Comprehensibility vs Immersivity: Rho=0.68; all p-values < 0.001).

**Table 2.** Perceptual ratings of the VR-AOT stimuli across five dimensions. Mean ratings of Comprehensibility, Realism, and Immersivity are presented for each of the 33 VR-based exercises, along with the perceived ownership reported from Egocentric and Allocentric perspectives. Each row corresponds to a different gesture, and each column presents the mean value across participants. Asterisks indicate a significant p-value at the one-sample t-test against 3 (p<0.05, Bonferroni corrected). NA indicates non-applicable scores for three exercises that cannot be presented from egocentric perspectives because their execution develops so close to the agent’s body to impede a proper first-person action observation.

|  | Scene Name | Comprehensibility | Realism | Immersivity | Egocentric | Allocentric |
| --- | --- | --- | --- | --- | --- | --- |
| E1 | Hit Ball - dorsal | 4,6* | 3,7* | 4,3* | 3,4 | 3 |
| E2 | Hit Ball - palmar | 4,7* | 4* | 4,4* | 4* | 3,4 |
| E3 | Hit Ball - Flex | 4,8* | 4,1* | 4,6* | 4,2* | 3,5 |
| E4 | HitBall - Ext | 4,4* | 3,8* | 4,6* | 4,1* | 3,5 |
| E5 | Drag soap | 4,6* | 4,3* | 4,6* | 4,2* | 3,5 |
| E6 | Drag ball | 4,5* | 4* | 4,5* | 4,1* | 3,5 |
| E7 | Drag cloth - flex | 4,8* | 4,4* | 4,7* | 4,2* | 3,5 |
| E8 | Reaching knee-table | 4,7* | 4,4* | 4,6* | 4,5* | 3,7 |
| E9 | Reaching a high target | 4,6* | 4,1* | 4,5* | 4,2* | 3,5 |
| E10 | Reaching lateral | 3,6* | 3,5 | 4,4* | 3,6* | 3,2 |
| E11 | Drag bottle | 4,8* | 4,4* | 4,6* | 4,3* | 3,6 |
| E12 | Move can | 4,6* | 3,9* | 4,6* | 4,1* | 3,5 |
| E13 | Bring soap | 4,4* | 3,7* | 4,4* | 3,9* | 3,1 |
| E14 | Open box | 4,6* | 4,1* | 4,6* | 4,1* | 3,3 |
| E15 | Drag box | 4,5* | 4* | 4,5* | 4* | 3,4 |
| E16 | Drag journal | 4,3* | 3,5 | 4,4* | 3,7* | 3,1 |
| E17 | Pour bottle | 4,8* | 4,4* | 4,6* | 4,3* | 3,6 |
| E18 | Clean table | 4,8* | 4,4* | 4,6* | 4,3* | 3,5 |
| E19 | Rolling pin | 4,2* | 3,2 | 4,4* | 3,6* | 3,2 |
| E20 | Flip glass | 4,1* | 3,4* | 4,4* | 3,7* | 3,2 |
| E21 | Press phone-key | 4,5* | 4* | 4,5* | 4* | 3,3 |
| E22 | Play piano | 4,6* | 4,2* | 4,4* | 4,1* | 3,5 |
| E23 | Hammer | 4,7* | 3,7* | 4,5* | 3,9* | 3,3 |
| E24 | Round spoon | 4,7* | 4* | 4,5* | 4* | 3,3 |
| E25 | Plug-unplug | 4,2* | 3,6* | 4,4* | 3,7* | 3,3 |
| E26 | Grasp cubes | 4,7* | 4,2* | 4,5* | 4,1* | 3,4 |
| E27 | Open jar | 4,1* | 2,7 | 4,3* | 3,1 | 2,8 |
| E28 | Handle | 4,2* | 3,3 | 4,3* | 3,6* | 3,2 |
| E29 | Writing | 4,1* | 3,2 | 4,4* | 3,5 | 3,1 |
| E30 | Sprayer | 4,8* | 4,3* | 4,5* | 4,2* | 3,5 |
| E31 | Throw ball - L | 4* | 3,3 | 3,5 | NA | 2,8 |
| E32 | Throw ball - H | 4,1* | 3,4 | 3,5 | NA | 2,7 |
| E33 | Toothbrush | 4,1* | 2,9 | 3,6 | NA | 2,9 |

Despite both egocentric (average 4.0 ± 0.3) and allocentric (average 3.3 ± 0.3) perspectives being rated positively in terms of ownership, a significant difference was found between them (Wilcoxon signed-rank test, W = 465, *p* < 0.001), suggesting that each perspective triggers the embodiment process differentially during action observation.

Overall, the results validate that the stimuli reported in the dataset preserved satisfactory perceptual quality throughout the kinematics-to-avatar rendering process, as evidenced by the ceiling scores for Comprehensibility and Realism of the movement. Combined with the immersive nature of the dataset and the enhanced embodiment elicited by the egocentric perspective, these VR-AOT stimuli show strong potential for use in VR-based motor rehabilitation protocols.

### Clinical utility of the dataset in stroke motor recovery

The Brunnström approach to stroke motor recovery outlines six stages, describing the progression of movement and coordination after a stroke. These stages are: flaccidity, spasticity appears, increased spasticity, decreased spasticity, complex movement combinations, and spasticity disappears. Considering this variety, it is highly unlikely that any exercise will be suitable for all of them. This assumption reflects pretty well in the stage-dependent course of the suitability scores for each exercise (see Fig. 3, Panel a). Overall, it is worth noting that most of the developed stimuli (except the Reaching lateral, Handle, and Throw ball - L and Throw ball - H) achieved a significantly positive score in at least one stage, demonstrating their suitability for the stroke motor rehabilitation framework. In addition, an increasing trend is observed in most exercises, indicating that the selected gesture repertoire may adapt more to later stages of motor recovery (IV to VI) than to its initial phases. However, some variance can be observed within our dataset, with half of the gestures already exploitable at stage IV, and the remaining ones turning significantly positive only at stages V and VI.

**Figure 3.**
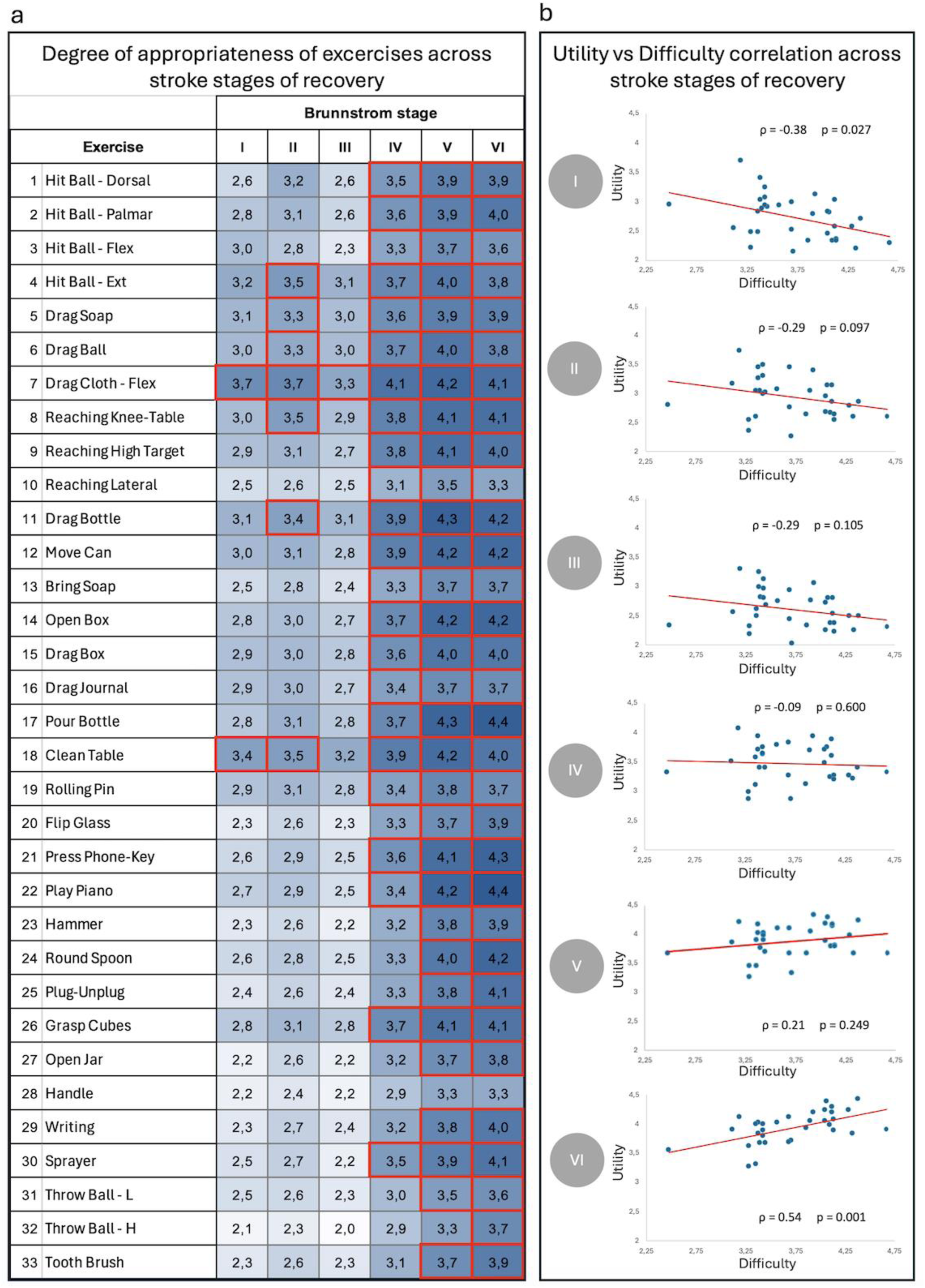
Clinical utility and its relationship with task difficulty across the Brunnström stroke recovery stages. Panel a: Mean utility ratings for each of the 33 VR gestures across the six Brunnström stages of stroke recovery (I–VI). Each cell color reflects the extent of the utility score, while red contours indicate scores significantly above the threshold of 3 (Wilcoxon signed-rank test, *p* < 0.05, Bonferroni-corrected), demonstrating the widespread perceived suitability of the gestures across all recovery stages. Panel b: Scatter plots display the relationship between the mean ratings of utility and difficulty, along the Brunnström stages of stroke recovery (I-VI). Each panel shows a linear regression line (red) and the Spearman’s correlation coefficient (ρ), along with its p-value.

This variance reflects that, even if our exercises align with the mid-to-late phase of motor recovery from stroke, their difficulty range makes them suitable for use at various stages of recovery. This assumption was further supported by correlation analyses examining the relationship between perceived utility for each Brunnström stage and task difficulty (Fig. 3 Panel b). Indeed, in the early phases of recovery (Stages I–III), correlations were weakly negative and close to the significance threshold (Stage I: ρ = –0.38, *p* = 0.027; Stage II: ρ = –0.29, *p* = 0.097; Stage III: ρ = – 0.29, *p* = 0.105), suggesting that simpler tasks were viewed as more useful and reflecting a prioritization of feasibility over challenge.

In contrast, the later stages (IV–VI) showed a shift toward positive associations between utility and difficulty. While the correlation in Stage IV is negligible (ρ = 0.09, *p* = 0.60), a moderate trend emerged at Stage V (ρ = 0.21, *p* = 0.249), culminating in a significant and substantially positive correlation at Stage VI (ρ = 0.54, *p* < 0.001). This pattern suggests that as functional recovery progresses, more complex tasks are perceived as increasingly valuable, aligning with the principles of progressive motor rehabilitation. Overall, these results highlight the adaptability of our VR-AOT dataset to various stages of motor recovery, underscoring its potential application in personalized rehabilitation pathways.

### Clinical utility of the dataset among other diagnostic domains

Finally, the stimuli were evaluated for their clinical utility for seven additional diagnostic domains drawing onto motor rehabilitation in their clinical course, namely Parkinson’s Disease (PD), Multiple Sclerosis (MS), Pediatric Cerebral Palsy (CP), Traumatic Brain Injury (TBI), Peripheral Nervous System pathologies (PNS), orthopedic trauma (OT), and prosthetic rehabilitation (PROSTH) (Table 3). Most stimuli obtained positive utility scores, except for the Reaching Lateral and Handle exercises, which did not surpass the three threshold in some conditions. Thus, the overall pattern suggests a broad perception of relevance across diverse neurological and musculoskeletal conditions. However, the results of the statistical analysis identify two distinct groups of clinical conditions. On one side, the highest utility is associated with clinical conditions (e.g., PNS, OT, prostheses) that primarily impact peripheral districts and spare the brain’s motor centers. More than half of the exercises were rated significantly positive for these conditions (20 for OT, 23 for PNS, 20 for PROSTH), with the three sets of exercises not completely overlapping, thus ruling out that exercises were rated solely based on their perceptual quality.

**Table 3.**
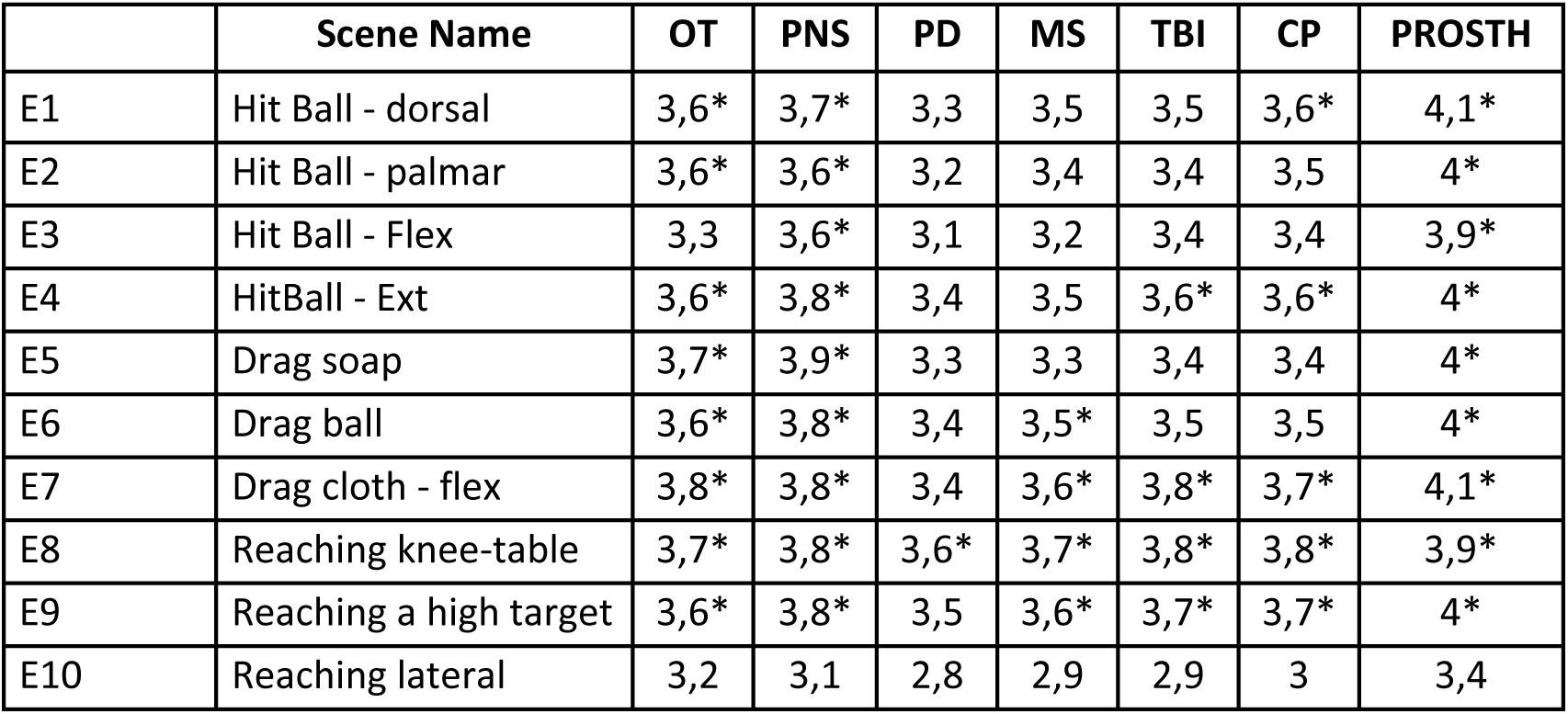
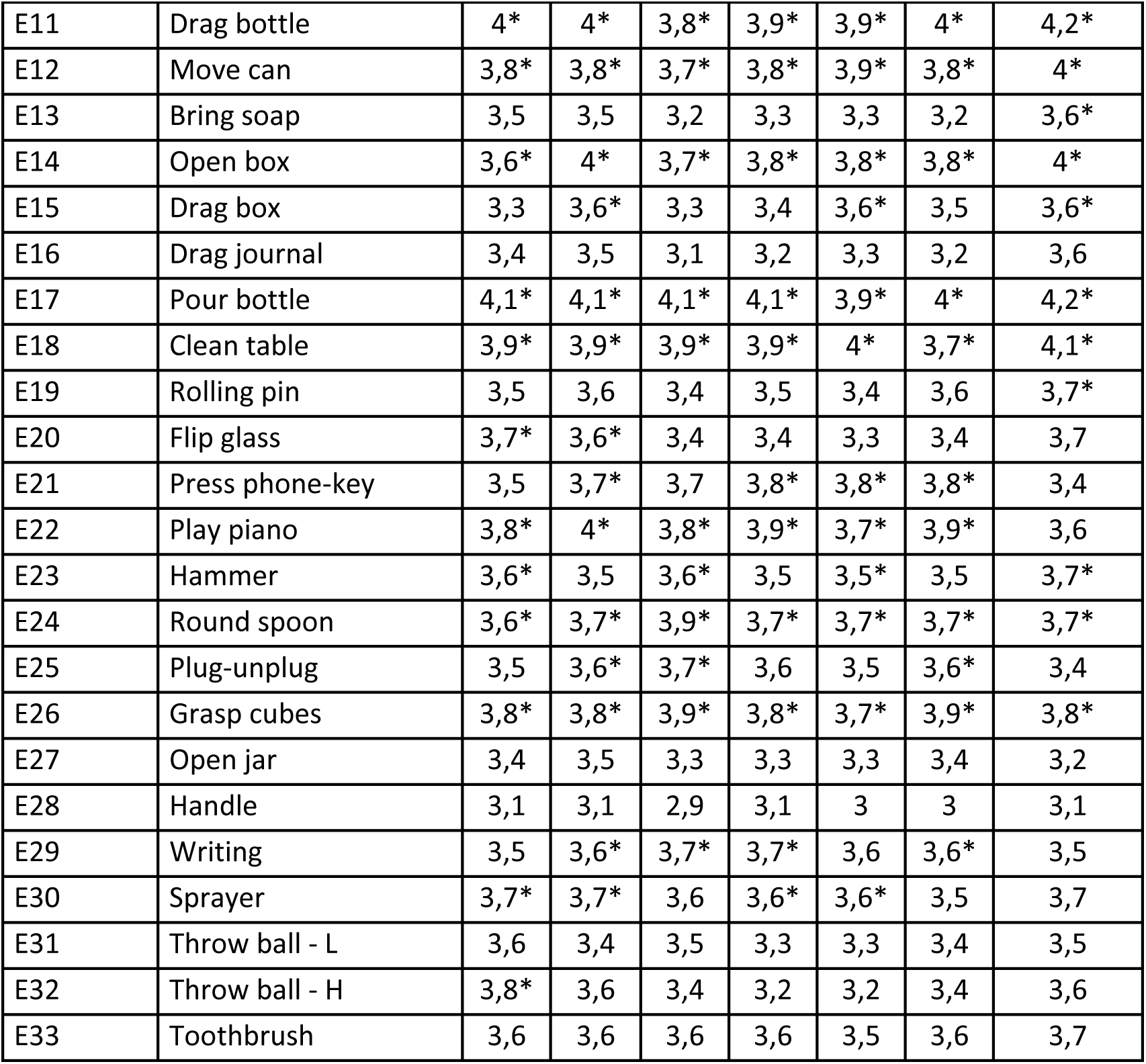
Suitability of VR-AOT exercises across neurological and musculoskeletal conditions. For each of 33 exercises, the degree of utility across different pathological conditions is reported. Asterisks indicate a significant p-value at the Wilcoxon signed-rank test against 3 (p<0.05, Bonferroni corrected). OT: Orthopaedic conditions; PNS: Peripheral Nervous System diseases; PD: Parkinson’s disease; MS: Multiple Sclerosis; TBI: Traumatic Brain Injury; CP: Cerebral Palsy; PROSTH: Prosthetic rehabilitation.

|  | Scene Name | OT | PNS | PD | MS | TBI | CP | PROSTH |
| --- | --- | --- | --- | --- | --- | --- | --- | --- |
| E1 | Hit Ball - dorsal | 3,6* | 3,7* | 3,3 | 3,5 | 3,5 | 3,6* | 4,1* |
| E2 | Hit Ball - palmar | 3,6* | 3,6* | 3,2 | 3,4 | 3,4 | 3,5 | 4* |
| E3 | Hit Ball - Flex | 3,3 | 3,6* | 3,1 | 3,2 | 3,4 | 3,4 | 3,9* |
| E4 | HitBall - Ext | 3,6* | 3,8* | 3,4 | 3,5 | 3,6* | 3,6* | 4* |
| E5 | Drag soap | 3,7* | 3,9* | 3,3 | 3,3 | 3,4 | 3,4 | 4* |
| E6 | Drag ball | 3,6* | 3,8* | 3,4 | 3,5* | 3,5 | 3,5 | 4* |
| E7 | Drag cloth - flex | 3,8* | 3,8* | 3,4 | 3,6* | 3,8* | 3,7* | 4,1* |
| E8 | Reaching knee-table | 3,7* | 3,8* | 3,6* | 3,7* | 3,8* | 3,8* | 3,9* |
| E9 | Reaching a high target | 3,6* | 3,8* | 3,5 | 3,6* | 3,7* | 3,7* | 4* |
| E10 | Reaching lateral | 3,2 | 3,1 | 2,8 | 2,9 | 2,9 | 3 | 3,4 |
| E11 | Drag bottle | 4* | 4* | 3,8* | 3,9* | 3,9* | 4* | 4,2* |
| E12 | Move can | 3,8* | 3,8* | 3,7* | 3,8* | 3,9* | 3,8* | 4* |
| E13 | Bring soap | 3,5 | 3,5 | 3,2 | 3,3 | 3,3 | 3,2 | 3,6* |
| E14 | Open box | 3,6* | 4* | 3,7* | 3,8* | 3,8* | 3,8* | 4* |
| E15 | Drag box | 3,3 | 3,6* | 3,3 | 3,4 | 3,6* | 3,5 | 3,6* |
| E16 | Drag journal | 3,4 | 3,5 | 3,1 | 3,2 | 3,3 | 3,2 | 3,6 |
| E17 | Pour bottle | 4,1* | 4,1* | 4,1* | 4,1* | 3,9* | 4* | 4,2* |
| E18 | Clean table | 3,9* | 3,9* | 3,9* | 3,9* | 4* | 3,7* | 4,1* |
| E19 | Rolling pin | 3,5 | 3,6 | 3,4 | 3,5 | 3,4 | 3,6 | 3,7* |
| E20 | Flip glass | 3,7* | 3,6* | 3,4 | 3,4 | 3,3 | 3,4 | 3,7 |
| E21 | Press phone-key | 3,5 | 3,7* | 3,7 | 3,8* | 3,8* | 3,8* | 3,4 |
| E22 | Play piano | 3,8* | 4* | 3,8* | 3,9* | 3,7* | 3,9* | 3,6 |
| E23 | Hammer | 3,6* | 3,5 | 3,6* | 3,5 | 3,5* | 3,5 | 3,7* |
| E24 | Round spoon | 3,6* | 3,7* | 3,9* | 3,7* | 3,7* | 3,7* | 3,7* |
| E25 | Plug-unplug | 3,5 | 3,6* | 3,7* | 3,6 | 3,5 | 3,6* | 3,4 |
| E26 | Grasp cubes | 3,8* | 3,8* | 3,9* | 3,8* | 3,7* | 3,9* | 3,8* |
| E27 | Open jar | 3,4 | 3,5 | 3,3 | 3,3 | 3,3 | 3,4 | 3,2 |
| E28 | Handle | 3,1 | 3,1 | 2,9 | 3,1 | 3 | 3 | 3,1 |
| E29 | Writing | 3,5 | 3,6* | 3,7* | 3,7* | 3,6 | 3,6* | 3,5 |
| E30 | Sprayer | 3,7* | 3,7* | 3,6 | 3,6* | 3,6* | 3,5 | 3,7 |
| E31 | Throw ball - L | 3,6 | 3,4 | 3,5 | 3,3 | 3,3 | 3,4 | 3,5 |
| E32 | Throw ball - H | 3,8* | 3,6 | 3,4 | 3,2 | 3,2 | 3,4 | 3,6 |
| E33 | Toothbrush | 3,6 | 3,6 | 3,6 | 3,6 | 3,5 | 3,6 | 3,7 |

On the other hand, neurological conditions primarily affecting the brain, and therefore also the motor system, received milder yet mostly positive ratings. CP and TBI patients were considered eligible for 16 exercises, MS patients for 15, and PD patients for 12 exercises. These findings underscore the dataset’s versatility and support its applicability across a range of neurorehabilitation contexts, including those beyond stroke recovery.

### Usage Notes

- **Editor compatibility.** It is crucial to open the project specifically with Unity version 6000.0.24f1 to ensure the compatibility and integrity of all scripts, prefabs, and associated assets. Using any other version, especially earlier releases, may result in the regeneration of asset identifiers, invalidation of prefab references, and potential incompatibilities that could impact reproducibility and stability.
- **Scene- level testing and export.** Each scene can be run and, if desired, exported as a standalone Unity package. Within the Unity Editor, you can enter Play Mode for a single gesture scene, adjust the desiredLoopCount parameter (found in the SceneDataManager component of the SceneManager GameObject, present in every scene) to control the number of repetitions the avatar performs, and interact with the game window using the mouse. When Meta Quest Link is active, the Game view is streamed directly to the headset, allowing real- time inspection in virtual reality.
- **Custom playback.** The project includes a dedicated scene, CustomScene, containing a GameObject named CustomManager. At runtime, you may supply an arbitrary, comma- separated list of scene IDs as a text string: the manager loads and plays them in the specified sequence, allowing repetitions of the same stimulus if required. An additional slider lets you select the inter- scene delay; during this interval, a black screen with a central fixation cross is presented. The feature is intended for fully customised stimulus delivery—e.g., experimental paradigms, demonstrations, or therapy sessions—and is available when running the project in the Unity Editor via **Meta Quest Link**.
- **APK deployment.** The UpperLimbAOT-VR.apk targets minSdkVersion 31 and the arm64- v8a ABI. Installation via the Meta Quest Developer Hub (drag- and- drop) is recommended. Ensure that ControlScene is designated as the launch scene.

These recommendations aim to streamline the user experience and facilitate the efficient incorporation of this dataset into diverse research workflows.

Any user should be aware that the current dataset is developed based on the kinematic recordings from a single healthy adult, which guarantees biomechanical consistency but may limit the generalizability of the kinematic profiles. In principle, we could have adopted normative kinematics^24^ to mitigate this issue; however, normative curves lose realism due to intra-population averaging, and reference datasets for upper-limb gestures, particularly those involving object interaction^25^, are fewer than gait-related ones and suffer from much larger interindividual variabilities.

Additionally, the repository currently focuses on unimanual gestures performed in a seated position. Although this design is typical of early to mid-stage rehabilitation, particularly in post-stroke recovery, it may not fully address the needs or tasks that require bimanual coordination and postural control in later stages of recovery. Future releases are planned to expand the database with bimanual and standing exercises, thus broadening its clinical applicability.

## Code Availability

Custom scripts were developed within the Unity engine (version 6000.0.24f1) to support both the Editor Mode and the Build Mode functionalities of the application. These scripts were specifically tailored to handle stimulus presentation, user interaction, data handling, and integration of motion capture animations. All scripts are fully documented and embedded within the Unity project that hosts the complete dataset.

## Acknowledgements

This work was supported by: the Italian Ministry of Health (Ricerca Finalizzata 2018, grant n. GR-2018-12367117) to PA; the Italian Ministry of Health (Ricerca Finalizzata 2021, grant n. RF-2021-12374941) to MAR and MFD; INAIL grant Virtualize (Piano Triennale della Ricerca, grant n. PR23-CR-P4) to PA and MFD; Fondazione Cariparma (BIOMONTANS project).

## Author contributions

**Ciraolo Alessandro**: Methodology, Software, Investigation, Formal Analysis, Writing-Original draft preparation, Writing- Reviewing and Editing.

**Scalona Emilia:** Methodology, Software, Writing- Original draft preparation, Writing-Reviewing and Editing.

**Zilli Adolfo:** Software

**Nuara Arturo:** Data Curation, Writing – Original Draft Preparation, Writing – Review & Editing

**De Marco Doriana:** Methodology, Data Curation

**Adamo Paola:** Conceptualization, Methodology, Writing – Review & Editing

**Gatti Roberto**: Conceptualization, Methodology, Writing – Review & Editing

**Rossi Paolo:** Data Curation, Writing – Review & Editing

**Banfi Sara:** Methodology, Data Curation

**Rocca Maria A:** Conceptualization, Funding Acquisition, Writing – Review & Editing

**Filippi Massimo:** Writing – Review & Editing

**Avanzini Pietro:** Conceptualization, Funding Acquisition, Supervision, Writing – Original Draft Preparation, Writing – Review & Editing

**Fabbri-Destro Maddalena:** Conceptualization, Funding Acquisition, Supervision, Writing – Original Draft Preparation, Writing – Review & Editing

